# PD-1 expression during acute infection is repressed through a LSD1- Blimp-1 axis

**DOI:** 10.1101/645838

**Authors:** Alexander P. R. Bally, Dennis K. Neeld, Peiyuan Lu, Parimal Majumder, Yan Tang, Benjamin G. Barwick, Qing Wang, Jeremy M. Boss

**Author notes:** These authors contributed equally to the work (Alexander P. R. Bally, Dennis K. Neeld, Peiyuan Lu). Corresponding Author: Jeremy M. Boss.

## Abstract

During prolonged exposure to antigens, such as chronic viral infections, sustained T cell receptor (TCR) signaling can result in T cell exhaustion mediated in part by expression of Programmed cell death-1 (PD-1) encoded by the *Pdcd1* gene. Here, dynamic changes in histone H3K4 modifications at the *Pdcd1* locus during ex vivo and in vivo activation of CD8 T cells, suggested a potential role for the histone H3 lysine 4 demethylase LSD1 in regulating PD-1 expression. CD8 T cells lacking LSD1 expressed higher levels of *Pdcd1* mRNA following ex vivo stimulation, as well as increased surface levels of PD-1 during acute but not chronic infection with lymphocytic choriomeningitis virus (LCMV). Blimp-1, a known repressor of PD-1, recruited LSD1 to the *Pdcd1* gene during acute but not chronic LCMV infection. Loss of DNA methylation at *Pdcd1*’s promoter proximal regulatory regions is highly correlated with its expression. However, following acute LCMV infection where PD-1 expression levels return to near base line, LSD1-deficient CD8 T cells failed to remethylate the *Pdcd1* locus to the levels of wild-type cells. Finally, in a murine melanoma model, the frequency of PD-1 expressing tumor infiltrating LSD1-deficient CD8 T cells was greater than wild-type. Thus, LSD1 is recruited to the *Pdcd1* locus by Blimp-1, downregulates PD-1 expression by facilitating the removal of activating histone marks, and is important for remethylation of the locus. Together, these data provide insight into the complex regulatory mechanisms governing T cell immunity and the regulation of a critical T cell checkpoint gene.

**Key Points:**

1. LSD1 suppress PD-1 expression following acute infection or transient induction.
2. Blimp-1 binding to the Pdcd1 locus is required to recruit LSD1.
3. LSD1 is required to fully remethylate the PD-1 proximal promoter region.

## Introduction

Programmed cell death-1 (PD-1) is an immunoinhibitory protein that is expressed on lymphocytes following the engagement of their antigen specific receptor (1). During the course of an acute infection, PD-1 expression is transient on the surface of CD8 T cells, peaking during the height of an infection and returning to near baseline levels in the resulting memory T cell pool (1–5). During chronic exposure to antigen, such as that from a chronic viral infection or in certain cancers, PD-1 expression is sustained at high levels and a state of T cell exhaustion is induced that is characterized by severe curtailment in effector functions, including the ability to proliferate, produce cytokines, and carry out cytotoxic responses (6). T cell exhaustion is in part mediated by signaling through PD-1’s intracellular tyrosine immunoinhibitory domains following PD-1’s surface engagement with its ligands PD-L1 or PD-L2 on target cells (1, 7–9). Although antibody blockade of PD-1/PD-L1 interactions can temporarily reinvigorate immune function from the exhausted state (1), PD-1 remains stably expressed on T cells during a chronic infection (10), and is expressed even upon removal of the cell from a chronic stimulatory environment (11). The stability of PD-1 expression and inhibition of effector functions across generations of cell division suggests that an epigenetic program stably regulates the transcriptional state of the *Pdcd1* locus.

PD-1 is encoded by the *Pdcd1* gene. In CD8 T cells, *Pdcd1* is regulated by the direct actions of transcription factors and epigenetic mechanisms (2, 10, 12). Upon T cell receptor engagement, *Pdcd1* is directly activated by a combination of transcription factors (NFATc1 and AP-1) that bind to a series of promoter proximal elements termed Conserved Regions (CR)-B and CR-C (12, 13), respectively. NFAT also binds to sites at −3.7 and +17.1 kb with respect to the transcription start site (14). Additional transcription factors appear to sustain *Pdcd1* during chronic infection and include the binding of NUR77 and FOXN1 to sites located at −23 kb and CR-C, respectively (15–18). Cytokine stimulation that results in STAT3 and STAT4 activation can further induce or sustain *Pdcd1* expression in mouse CD8 T cells by binding to the distal −3.7 and +17.1 sites (14). Following the cessation of the TCR signaling (e.g., via antigen/viral clearance), *Pdcd1* is silenced through the binding of B lymphocyte induced maturation protein-1 (Blimp-1) to a region between CR-C and CR-B (19). Blimp-1 binding results in the eviction of NFATc1 from CR-C (19). In addition to these mechanisms, epigenetic regulation through DNA methylation occurs across CR-C and near CR-B in CD8 T cells in both mice and humans (2, 10). In naïve CD8 T cells, CpGs in the above DNA regions are consistently methylated. Upon CD8 T cell activation, the methylation is lost in a time course that parallels *Pdcd1* expression. In an acute infection setting, the locus is remethylated as the infection is cleared and PD-1 levels return to the baseline as mentioned above. By contrast, during chronic infection, DNA methylation is permanently lost and is not regained (2, 10).

In a similar manner, the accumulation and removal of activating and repressing histone modifications correlate completely with *Pdcd1* expression in mouse CD 8 T cells. For example, both H3K27^ac^ and H3K9^ac^ activation modifications at CR-B and CR-C correlate with *Pdcd1* expression when driven by ex vivo TCR stimulation (19, 20), and H3K4^me1^ is enriched when the above stimulation is coupled with IL-6 and IL-12 (STAT3/STAT4) treatment (14). However, as expression wanes during ex vivo stimulation, the repressive modifications H3K9^me3^, H3K27^me3^, and H4K20^me3^ appear at CR-B and CR-C (19). Using the EL4 T cell line, exogenous expression of the transcriptional repressor Blimp-1 induced the appearance of all three of these repressive modifications at the *Pdcd1* locus and subsequently silenced PD-1 expression (19). Surprisingly, Blimp-1 is also expressed during a chronic infection in exhausted CD8 T cells where PD-1 levels are at their highest, yet fails to repress PD-1 (21). The molecular mechanism for how Blimp-1 could function to repress *Pdcd1* exclusively following acute inflammation is not fully clear.

As a repressor, Blimp-1 is known to recruit additional transcriptional repressors that result in silencing the local chromatin environment (22–24). Along with histone deacetylases HDAC1 and HDAC2 and the histone methyltransferase G9a, Blimp-1 can recruit the lysine-specific demethylase 1a (LSD1) encoded by *Kdm1a* (23–25). LSD1 catalyzes the removal of mono and dimethylation modifications of H3K4 that are associated with transcriptional activation (26–28), thereby facilitating an epigenetic state of gene silencing. Given the ability of Blimp-1 to repress genes through recruitment of histone modifiers such as LSD1, we set out to test the hypothesis that LSD1 contributes to the regulation of *Pdcd1* in a Blimp-1-dependent manner. Indeed, we found that Blimp-1 was necessary to recruit LSD1 to the *Pdcd1* locus, and that when bound, LSD1 actively down-regulates PD-1 transcription and expression. Furthermore, we found that Blimp-1 was bound to the *Pdcd1* locus in both acute and chronic settings; however, LSD1 was only recruited following an acute stimulation, correlating with removal of proximal H3K4^me1/me2^ modifications and appearance of a repressive epigenetic profile concurrent with *Pdcd1* silencing and DNA remethylation of the locus following acute viral infection. We also found that greater frequency of PD-1 expressing tumor infiltrating CD8 T cells in LSD1-deficient compared to LSD1-sufficient mice in a melanoma model. Thus, LSD1 and Blimp1 together are responsible for resetting the epigenetic programming of the *Pdcd1* locus to a resting state.

## Material and Methods

### Animals

C57BL/6J mice were obtained from Jackson laboratories. *Kdm1a*^fl/fl^ mice (29) (provided by D. Katz at Emory University) were crossed to mice containing a Granzyme B promoter-driven Cre recombinase (B6;FVB-Tg(GMB-cre)1Jcb) (provided by J. Jacob at Emory University) (30), and subsequently back-crossed to the C57BL/6 mouse line for 4-5 generations. *Prdm1*^fl/fl^ mice were provided by K. Calame (Columbia University) and similarly crossed to Granzyme B-cre mice (19). Equal numbers of male and female mice were used in all experiments. The numbers of animals used in each experiment are provided in the figure legends. All experiments were performed in accordance with approved protocols by the Emory University Institutional Animal Care and Use Committee.

### Virus Infection

Viral stocks of lymphocytic choriomeningitis virus (LCMV) strains Armstrong and Clone-13 were generated as previously described (31) and kindly provided by Dr. Rafi Ahmed (Emory University). Mice were infected with 2×10^5^ pfu of LCMV Armstrong intraperitoneally or with 2×10^6^ pfu LCMV Clone-13 intravenously as described (32). Viral titers were determined by plaque assay using Vero cells (ATCC CCL-81) as previously described (32, 33)

### Cell isolation and ex vivo cell activation

For analysis or ex vivo cell culture, CD8 T cells were isolated from single-cell splenocyte suspensions using the Miltenyi CD8a+ T cell isolation kit (Miltenyi Biotec, Inc. Cat.130-104-075) according the manufacture’s protocol. Where indicated, LCMV tetramer specific cells (see below) were further purified by FACS on a BD FACSAria II (Emory SOM Flow Cytometry Core) following infection as indicated and biochemically/molecularly analyzed immediately. For some experiments, isolated cells were cultured ex vivo in RPMI supplemented with 5% FBS, 5% bovine calf serum, 4.5 g/l glucose, 1.0 mM sodium pyruvate, 10 mM HEPES and 100 U/ml Penicillin/streptomycin. For ex vivo activation, isolated CD8 T cells were stimulated using Dynabeads Mouse T-activator CD3/CD28 kit (Thermo Fisher Scientific (Gibco) Cat. 11453D) according to the manufacture’s protocol at a ratio 2:1 beads/cell for the indicated time (24-96 hours).

### Flow cytometry

Cells were stained for flow cytometry in FACS buffer (PBS, 1% BSA, 1 mM EDTA) for 30 m, and subsequently fixed using 1% paraformaldehyde for 30 m. Events were collected on a BD LSR II and analyzed using FloJo 9 software. Antibodies used to stain cells included: CD4 PerCP-Cy5.5 (clone RM4.5); CD8 FITC (clone 53-6.7); CD44 APC-Cy7 (clone IM7); CD62L Alexa Fluor 700 (clone MEL-14); CD69 PE-Cy7 (Clone H1.2F3); CD127 BV510 (Clone SB/199); PD-1 PE (clone RMP1-30). Biotinylated H-2D^b^ MHC tetramers specific for LCMV peptides for gp33 var C41M (KAVYNFATM), gp276 (SGVENPGGYCL), and np396 (FQPQNGQFI) were obtained from the NIH Tetramer Core facility at Emory University and subsequently tetramerized to streptavidin-APC (Prozyme) following their protocols (tetramers.yerkes.emory.edu).

### Quantitative Real-time PCR

RNA was isolated from at least three independent preparations of cells using the RNAeasy kit (Qiagen, Inc.), and cDNA was prepared from RNA libraries using SuperScript II reverse transcriptase (Life Technologies Co). RT-PCR was used to quantitate mRNA levels in technical duplicates, and values were normalized using 18s ribosomal RNA as previously described (34).

### Chromatin Immunoprecipitation

Chromatin immunoprecipitation (ChIP) assays were performed as previously described (19, 35). Briefly, purified cell populations were crosslinked for 10 minutes in 1% formaldehyde, and subsequently lysed in cell lysis buffer (5mM PIPES pH 8.0, 85 mM KCl, 0.5% NP-40). Chromatin was extracted using nuclei lysis buffer (50 mM TRIS pH 8.1, 10 mM EDTA, 1% SDS) and sonicated to an average length of 400-600 bp. Chromatin (5 μg) was used for immunoprecipitation reactions on Protein A beads with 0.5 μg of polyclonal antibodies for H3K4^me1^ (Millipore Sigma Cat. 07-436), H3K4^me2^ (Millipore Sigma Cat. 07-030), H3K4^me3^ (Millipore Sigma Cat. 07-473), H3K27^ac^ (Millipore Sigma. Cat. 07-360), IgG (Millipore Sigma, Cat. 12-370), Blimp-1 (Rockland Cat. 600-401-B52) and LSD1 (Santa Cruz Cat. SC-271720). Precipitates were quantitated by quantitative PCR and calculated as a percent of input. (Supplemental Figure 1).

### DNA methylation analysis

The DNA methylation content of the CR-B associated region was determined by clonal bisulfite sequencing as previously described (2). Briefly, genomic DNA purified from CD8 T cells and bisulfite converted using the EpiTect Bisulfite Kit as per the manufacturer’s instructions (Qiagen, Inc.). Bisulfite-converted DNA was PCR amplified and cloned with the TOPO TA cloning kit (Life Technologies). Clones were isolated and the plasmid DNA regions were sequenced and changes in CpG DNA methylation were determined. Data were aligned *in silico* using the R / Bioconductor Biostrings package and custom scripts as previously described (36). A Fisher exact test was used to determine significance.

## Results

### Pdcd1 *histone modifications differ between acute and chronic infection*

As previously shown (1, 2), infection with LCMV Armstrong induces an acute infection that results in CD8 T cells expressing high surface levels of PD-1 on day 5 post-infection, but this ultimately decreases to near naïve T cell levels 8 days following infection when virus is no longer detectable (Fig.1A). Conversely, infection with the chronic LCMV strain Clone-13 results in PD-1 surface (Fig. 1A) and mRNA levels (2) that remain elevated during the course of the infection. To gain additional insight into epigenetic mechanisms that may be driving the above changes and differences in PD-1 expression between acute and chronic infections, we examined activating and repressive histone modifications at key regulatory elements (37) across the *Pdcd1* locus (Fig. 1B). At day 8, splenic CD8 T cells were isolated and analyzed by chromatin immunoprecipitation (ChIP). We observed that the −3.7 and 17.1 regions were enriched for the active H3K4 mono-methylated histone mark (H3K4^me1^) in antigen specific CD8 T cells of chronically infected mice compared to both acutely infected and naïve mice (Fig. 1C). Another active histone mark, H3K4^me2^, was also enriched in chronically infected mice at both the CR-B and CR-C regulatory regions of the *Pdcd1* locus. H3K27^ac^ enrichment, a mark associated with active promoters/enhancer regions (38) was higher at all sites in CD8 T cells from Clone-13 infected mice compared to those from Armstrong infected or naïve mice. ChIP for the repressive histone marks H3K9^me2^ and H3K27^me3^ were enriched at both CR-B and CR-C but only in Armstrong infected mice (Fig.1C). Taken together, these data demonstrate that at day 8 post infection, CD8 T cells from chronically infected mice have active histone marks in the *Pdcd1* locus whereas the acutely infected mice have an enrichment for repressive marks. Moreover, these histone modifications correspond with PD-1 expression on CD8 T cells from chronically and acutely infected mice (Fig. 1A, C).

**Figure 1.**
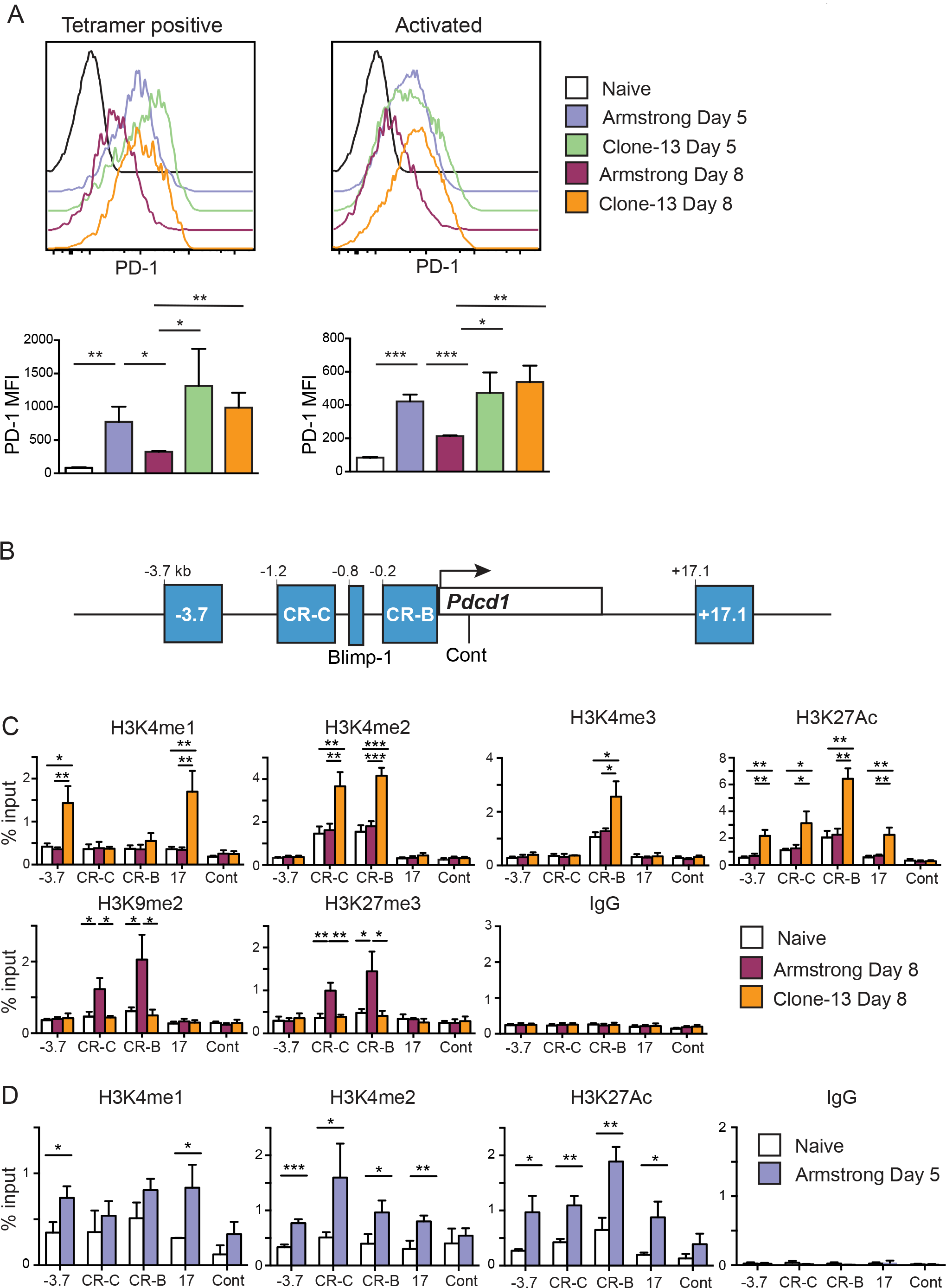
Activating histone marks are dynamically regulated and correlate with Pdcd1 expression. C57Bl/6 mice were infected with either LCMV Armstrong or Clone-13 for 5 or 8 days. (**A**) Representative flow cytometry plots showing the surface PD-1 levels on CD8 T cells. Plots were gated on CD8 T cells and show combined LCMV gp33, gp276 and np396 tetramer positive cells (day 8) or total activated CD8 T cells defined as CD44^hi^ and CD62L ^low^ (day 5). Bar graphs represent combined averages of 3 mice. (**B**) Schematic highlighting regulatory elements of the *Pdcd1* locus where ChIP assays were performed. (**C**) ChIP assay for both activating and repressive histone marks on CD8 T cells isolated from day 8 Armstrong or Clone-13 infected mice. (**D**) ChIP on activated CD44^hi^ CD62L ^low^ isolated from day 5 infected Armstrong mice and naïve uninfected mice for activating histone marks. Data represent three independently isolated biological replicates. Two tailed Student’s *t* tests were used to determine significance in these experiments. *, P < 0.05 **, P < 0.01 ***, P < 0.001.

The above data raise the question as to whether active histone marks examined appear at an earlier timepoint during an *in vivo* response to an Armstrong infection that would correlate with the increased surface expression. To test this, cells were isolated at day 5 following LCMV Armstrong infection, a time point in which PD-1 expression was high (Fig. 1A) and enough activated CD8 T cells (CD44^hi^ CD62L^low^) could be isolated for ChIP. In these experiments, H3K4^me2^ and H3K27^ac^ were enriched over control naïve CD8 T cells at all four key regulatory elements; whereas H3K4me1 was only significantly enriched at −3.7 and 17.1 (Fig 1D). Thus, active histone modifications at key regulatory elements are associated with PD-1 expression and during the course of an acute infection, these marks are replaced by repressive histone modifications as expression is silenced. This raises the question of how the dynamics of histone modifications at the *Pdcd1* locus are regulated.

### LSD1 represses PD-1 following ex vivo stimulation

LSD1 encoded by *Kdm1a* is responsible for catalyzing the removal of both H3K4^me2^ and H3K4^me1^ marks to an unmethylated state, a process that has been dubbed enhancer decommissioning as it can lead to gene silencing (26, 27). To test the hypothesis that LSD1 is actively recruited and responsible for down regulation of *Pdcd1* gene expression following an acute infection, we crossed an *Kdm1a*^*fl/fl*^ mouse (29) with the *Cre*^*GranzB*^ mouse (30). The *Kdm1a*^*fl/fl*^ allele allows for *Cre*-mediated deletion of LSD1’s amine oxidase catalytic domain (29). The *Cre*^*GranzB*^ allele is induced in CD8 T cells upon T cell activation (19, 30) leading to a conditional deletion of LSD1 in activated CD8 T cells. Using the *Kdm1a*^*fl/fl*^*Cre*^*GranzB*^ mouse (designated LcKO henceforth), we first determined whether an LSD1 deficiency could alter the kinetics of *Pdcd1* expression in an *ex vivo* activation system. Here splenic CD8 T cells from littermate *Kdm1a*^*fl/fl*^*Cre*^−^ controls (termed WT) and LcKO mice were purified and stimulated in culture using αCD3/CD28 beads over a 4-day period and expression of *Pdcd1* mRNA was measured by qRT-PCR at daily intervals. CD8 T cells from LcKO animals displayed significantly higher levels of *Pdcd1* mRNA compared to WT littermate controls at 24 hours, the peak of *Pdcd1* expression by this method of stimulation (Fig. 2A). Moreover, whereas WT *Pdcd1* mRNA levels decayed over the 96 h time course, *Pdcd1* mRNA levels remained significantly higher in the LcKO CD8 T cells at all time points after initial stimulation. Additionally, qRT-PCR for *Kdm1a* mRNA levels showed that the LcKO animals had a significant reduction in LSD1 mRNA compared to WT controls, confirming efficient deletion of the floxed *Kdm1a* alleles following stimulation (Fig. 2B). The lag phase prior to complete deletion of *Kdm1a* may account for the slight, although significantly less than in WT, decrease in *Pdcd1* expression after peak. These results demonstrate that LSD1 acts as a negative regulator of PD-1 expression in CD8 T cells following ex vivo stimulation.

**Figure 2.**
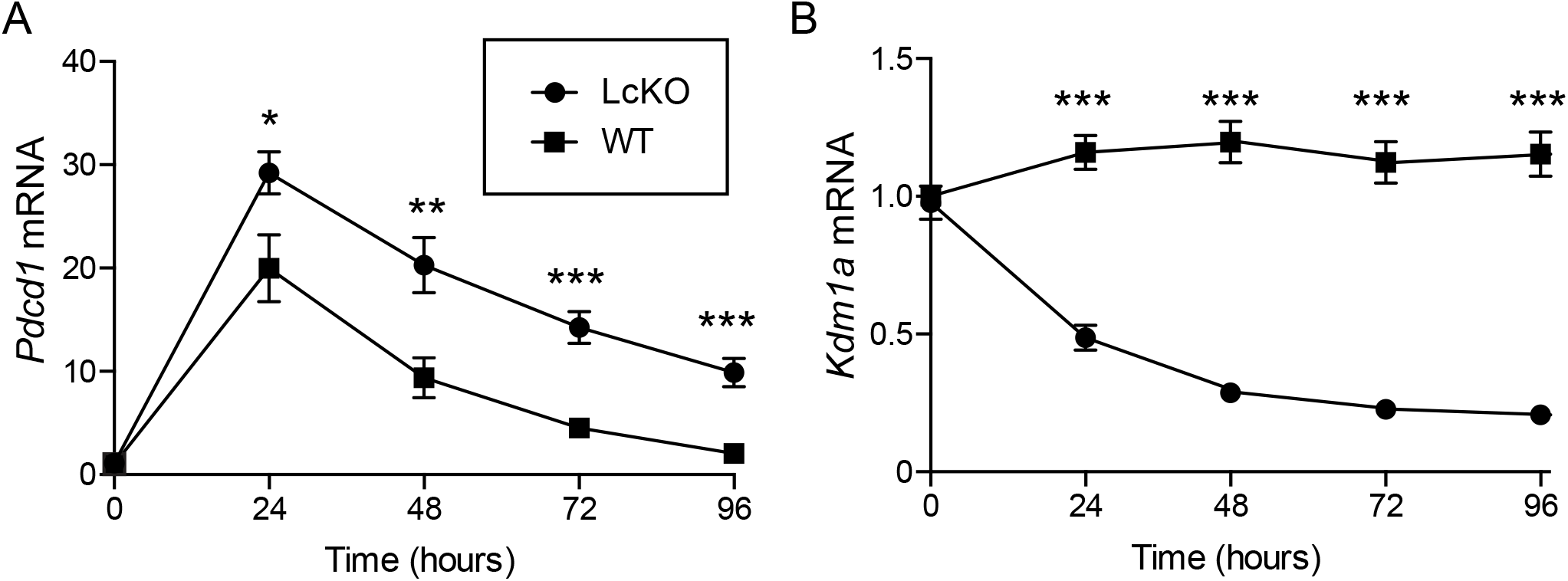
LSD1 acts as a repressor of *Pdcd1* expression in *ex vivo* stimulated CD8 T cells. CD8 T cells were isolated from *Kdm1a*^*fl/f*^*Cre*^−^ (WT) and *Kdm1a*^*fl/f*^*Cre*^*GranzB*^ (LcKO) by magnetic separation and then stimulated for up to 96 hours using αCD3/CD28 beads at a 2:1 bead:cell ratio. Every 24 hours, cells were collected for RNA and analyzed by qRT-PCR. (**A**) qRT-PCR showing *Pdcd1* expression. (**B**) qRT-PCR showing *Kdm1a* (LSD1) expression and that it was efficiently deleted from LcKO CD8 T cells. Data represent three independently isolated biological replicates. A two tailed Student’s *t* test was used to determine significance at each time point. *, P < 0.05 **, P < 0.01 ***, P < 0.001.

### LSD1 represses PD-1 following acute viral infection

To test if PD-1 expression is modulated by LSD1 during an acute viral infection, WT and LcKO mice were infected with LCMV Armstrong and PD-1 expression on LCMV antigen-specific CD8 T cells was monitored by flow cytometry. At day 8 post infection, when viral loads have been cleared and WT mice have down-regulated surface PD-1 expression on LCMV antigen-specific CD8 T cells (Fig. 3A, blue), LcKO CD8 T cells retained high levels of PD-1 expression (Fig. 3A red). Previously (19), we found that conditional deletion of Blimp-1, a transcriptional repressor that has been suggested to interact with LSD1 (23), produced a similar phenotype to LcKO mice, which is reconfirmed here (Fig. 3A) using *Cre*^*GranzB*^*Prdm1*^*fl/fl*^ (**BcKO**) mice. The similar phenotype suggests that these enzymes may operate along the same pathway, in line with the hypothesis that Blimp-1 recruits LSD1 in order to mediate *Pdcd1* repression. By 28 days post infection with Armstrong, LSD1 knockout antigen-specific CD8 T cells no longer showed an increase in PD-1 expression, unlike cells from Blimp-1 knockout mice, which still showed some level of PD-1 expression over their littermate (*Cre*^−^) controls (Fig. 3A). This suggests that while LSD1 is an important determinant in the pathway responsible for decreasing PD-1 expression it does not account for the entire level of PD-1 repression mediated by Blimp-1.

**Figure 3.**
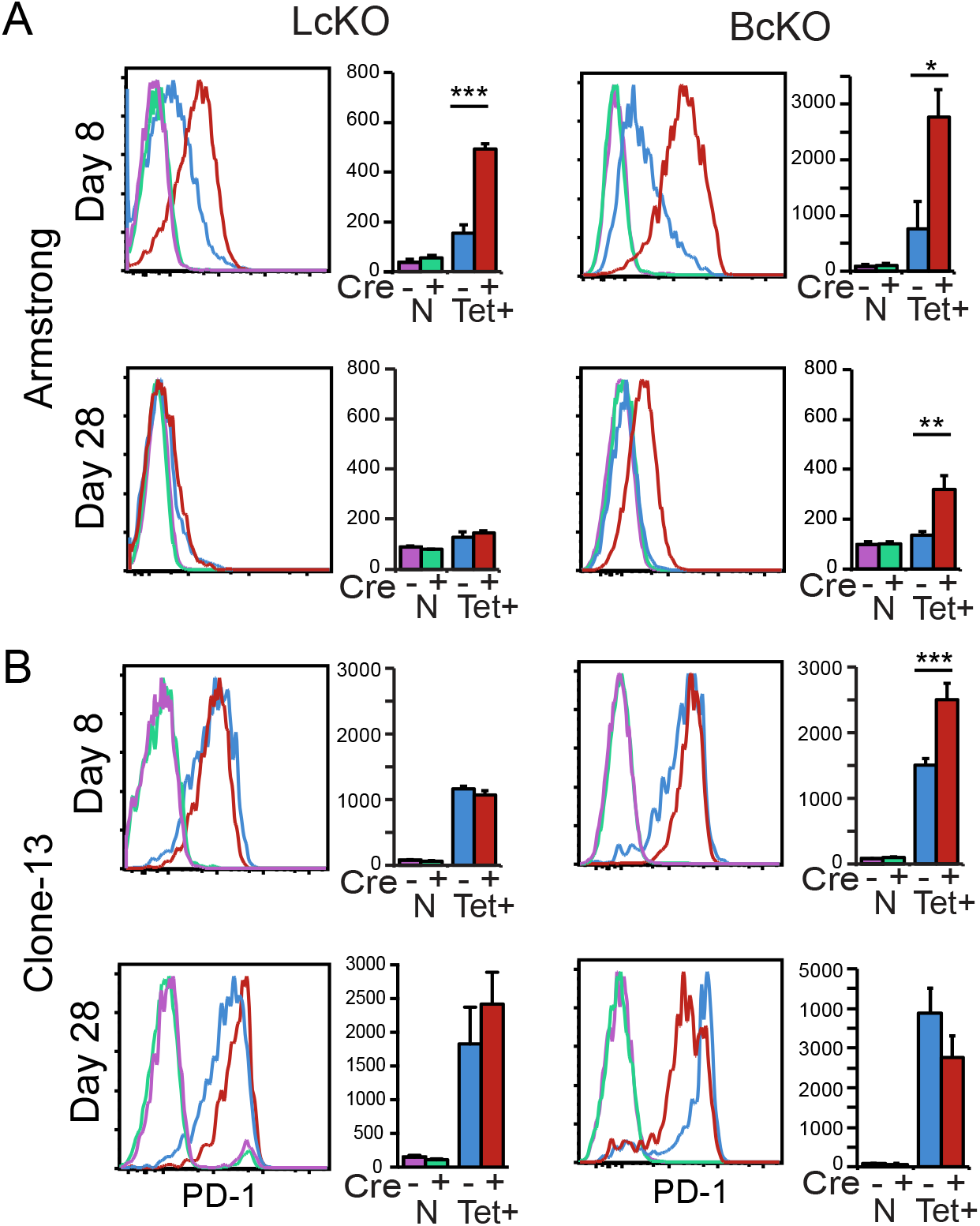
LSD1 represses PD1 during acute viral infection. LcKO, BcKO and WT control mice were infected with LCMV Armstrong or Clone-13 for 8 or 28 days. Cells were isolated and subject to flow cytometry to assay surface PD-1 expression. Plots show naïve (N) CD8 T cells gated as CD8^+^CD44^low^ CD62L^hi^; and LCMV-specific cells gated as CD8^+^, CD44^hi^ and gp33 tetramer positive (Tet+). Data represent three independently isolated biological replicates. A two tailed Student’s *t* test was used to determine the significance at each time point. *, P < 0.05 **, P < 0.01 ***, P < 0.001.

To determine if LSD1 affected PD-1 expression during a chronic LCMV infection, antigen specific cells were isolated from WT and LcKO mice following LCMV Clone-13 infection and analyzed by flow cytometry. At day 8, WT and LcKO CD8 T cells expressed moderate levels of PD-1 (Fig. 3B), while at day 28 post infection, no difference in MFIs were observed between WT and LcKO (Fig. 3, bottom). During chronic infection, BcKO CD8 T cells at day 8 had an overall mean fluorescent intensity that was higher, but the histogram patterns overlap with WT cells to a significant degree. Thus, at this early stage, Blimp-1 may not be contributing to *Pdcd1* regulation. At the chronic infection time point (day 28), BcKO CD8 T cells expressed slightly lower levels of PD-1, which was in agreement with previous reports (21). The differential roles performed by Blimp-1 and LSD1 at day 28 of a chronic infection may indicate that their activities may be dissociated under these conditions and at these late time points.

### *Blimp-1 recruits LSD1 to* Pdcd1

To determine if LSD1 interacts directly with the *Pdcd1* locus concurrent with PD-1 down-regulation, and whether it is recruited there by Blimp-1, ChIP was performed on *ex vivo α*CD3/CD28 stimulated splenic CD8 T cells isolated from LcKO and BcKO animals, as well as the corresponding *Cre*^−^ controls (WT). At day 4, when PD-1 expression has nearly returned to baseline levels (Fig. 2A), LSD1 was bound to the CR-B region in WT mice (Fig. 4A). This indicates a direct interaction with the locus and suggests that LSD1 itself directly represses *Pdcd1* transcription. However, LSD-1 binding was not observed in BcKO mice concurrent with a loss of Blimp-1 at the *Pdcd1* locus (Fig. 4A, B), indicating that Blimp-1 is necessary to recruit LSD1 to the region. Importantly, LSD1 binding was minimally observed at the locus in LcKO cells, indicating that the conditional deletion was efficient (Fig. 4A). However, Blimp-1 was still bound in LcKO mice (Fig. 4B), indicating that Blimp-1 is capable of independently interacting with *Pdcd1*. These data demonstrate that LSD1 interacts directly with the *Pdcd1* locus, corresponding with suppression of *Pdcd1* expression *ex vivo*, and requires Blimp-1.

**Figure 4.**
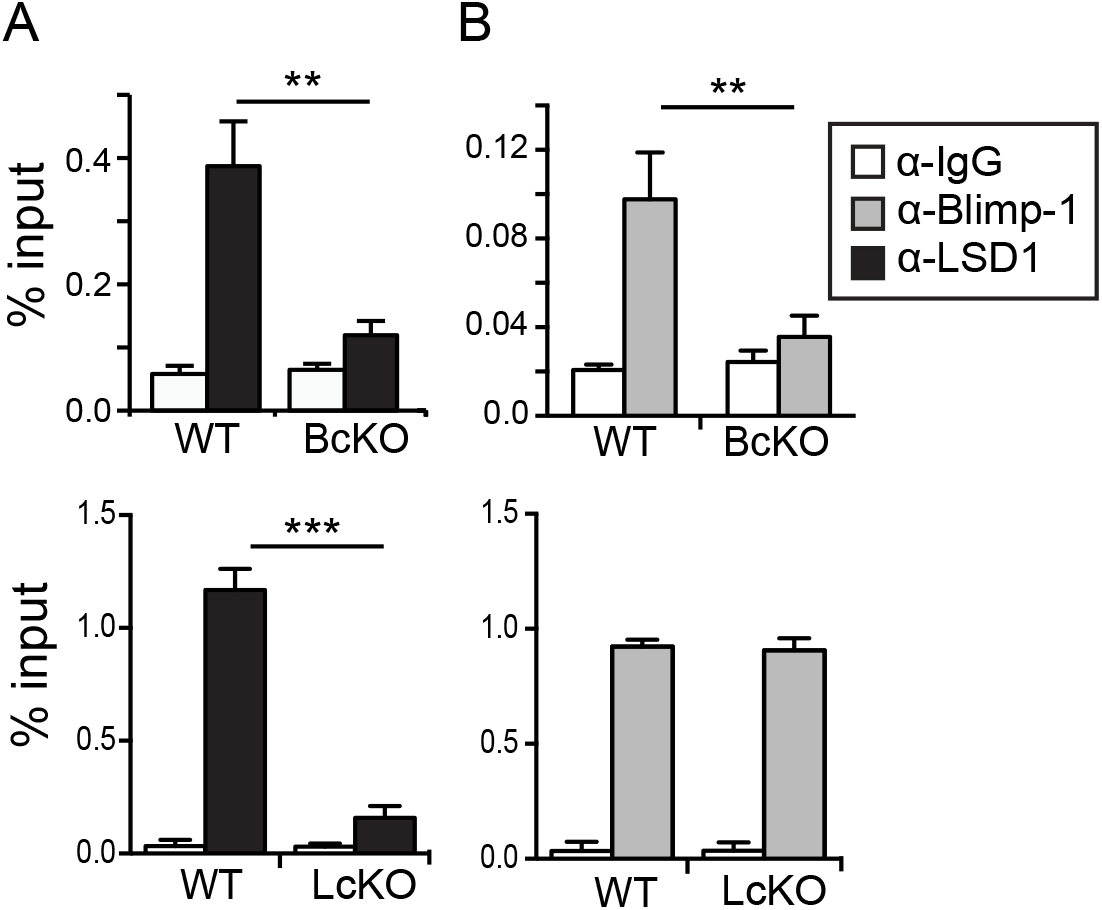
Blimp-1 recruits LSD1 to the *Pdcd1* locus. CD8 T cells from LcKO, BcKO, and WT control mice were isolated by magnetic separation and then stimulated with αCD3/CD28 beads for 96 hours. Chromatin was prepared from collected cells and then subject to ChIP. (**A**) Binding of LSD1 to the CR-B region of the *Pdcd1* locus. (**B**) Binding of Blimp-1 to site 2 of the *Pdcd1* locus. An α-IgG antibody was used as a control for ChIP experiments. Data represent three independently isolated biological replicates. A two tailed Student’s *t* test was used to determine statistical significance. **, P < 0.01 ***, P < 0.001.

### LSD1 interacts with Pdcd1 following an acute but not chronic infection

Blimp-1 binds directly to *Pdcd1* following an acute infection, and acts as a transcriptional repressor (19). However, during a chronic infection, Blimp-1 is expressed at even higher levels, and was correlated with maximal PD-1 expression (21). To determine if Blimp-1 interacted directly with *Pdcd1* during chronic inflammation *in vivo*, ChIP was performed on virus-specific CD8 T cells from naïve, day 8 acutely-infected (Armstrong), or day 8 chronically-infected (Clone-13) mice. As we previously reported, Blimp-1 was bound to *Pdcd1* following acute infection (19). LSD1 was also bound, correlating with its repressive role (Fig. 5A). Blimp-1 was also bound at day 8 during a chronic infection (Fig. 5A). Importantly, in this setting, LSD1 was not recruited to the region even though Blimp-1 was bound. To rule out the possibility of differential expression levels of LSD-1 between viral infections, we analyzed previously published (39) RNA microarray data and observed no differences (Fig. 5B). This suggests that although Blimp-1 interacts with the *Pdcd1* locus in CD8 T cells during a chronic infection setting, it fails to down-regulate PD-1 levels because LSD1 is not recruited.

**Figure 5.**
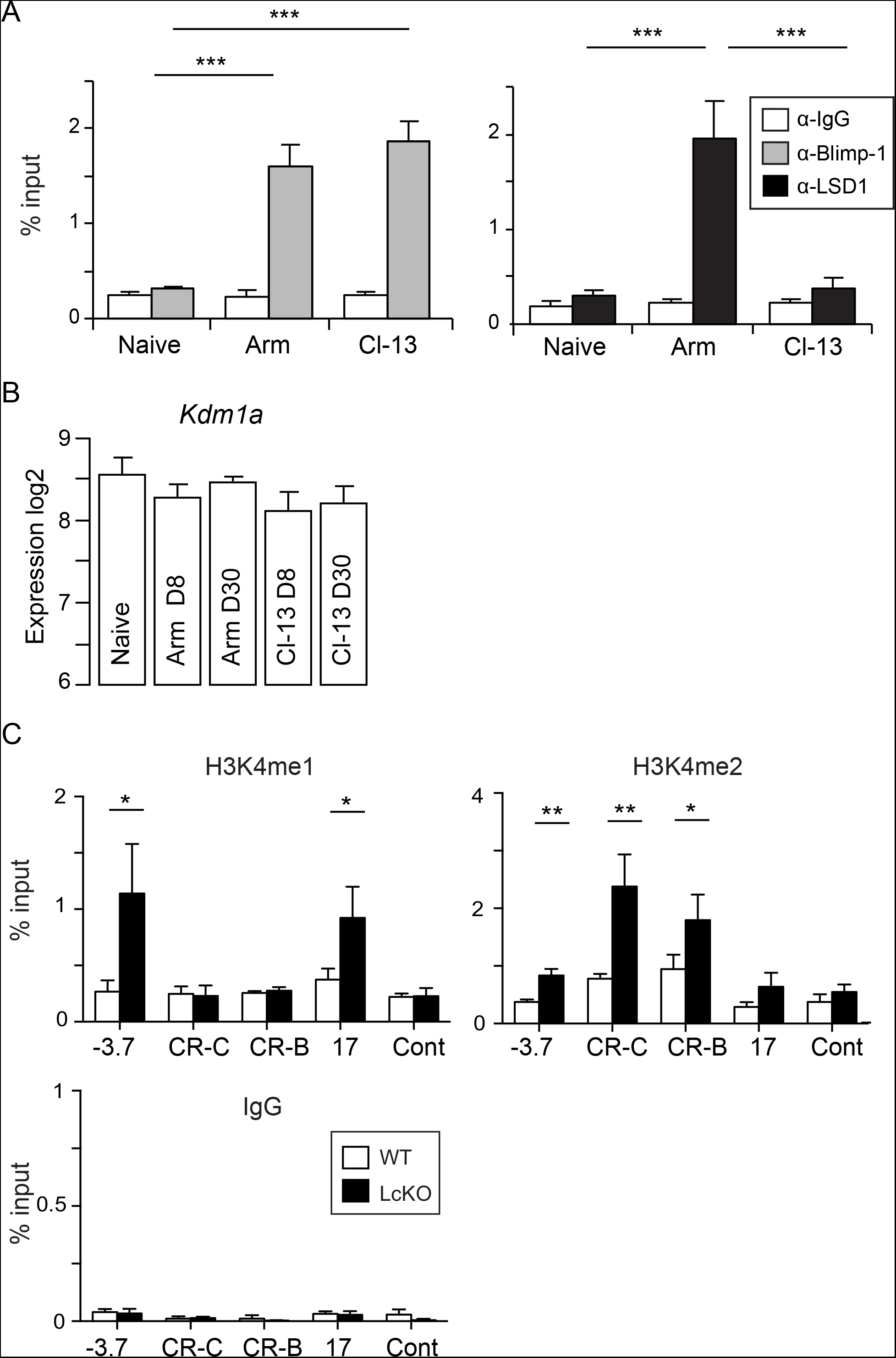
LSD1 is recruited to the *Pdcd1* locus only during an acute infection. (**A**) C57BL/6 mice were infected with either LCMV Armstrong or Clone-13 for 8 days. CD8 T cells were isolated by magnetic separation and subject to ChIP to assess LSD1 and Blimp-1 binding. Naïve CD8 T cells were also magnetically isolated (See Methods) from uninfected mice were used as a control, as was an α-IgG antibody. Blimp-1 binding was assessed at the Blimp-1 binding site (Fig 1B) of the *Pdcd1* locus and LSD1 binding was assessed at CR-B. (B) Microarray RNA expression data from (39) was analyzed for *Kdm1a* expression (LSD1). (**C**) LcKO and littermate control WT mice were infected with LCMV Armstrong. At day 8, gp33-, gp276-, and np396-specific CD8 T cells were isolated, pooled, and subjected to ChIP with α-H3K4^me1^ and H3K4^me2^ antibodies at the indicated regions of the *Pdcd1* locus. Three independent biological replicates were assayed. A two tailed Student’s *t* test was used to determine statistical significance between samples. *, P < 0.05 ***, P < 0.001.

### LSD1 is responsible for the decommissioning of Pdcd1 enhancer elements

As LSD1 is responsible for removing the histone marks H3K4^me1^ and H3K4^me2^, we set out to determine if LcKO mice possessed increased levels of these marks at the *Pdcd1* locus. To test this, LcKO and *Cre*^−^ control (WT) mice were infected with LCMV Armstrong for 8 days. Antigen specific (pooled gp33, gp276, np396) CD8 T cells were isolated by FACS and subjected to ChIP for H3K4^me1^ and H3K4^me2^ at key regulatory regions across the *Pdcd1* locus. CD8 T cells from LcKO mice showed enrichment for H3K4^me1^ at the −3.7 and +17.1 regions compared to WT cells (Fig. 5C). Enrichment for H3K4^me2^ was also observed in the LcKO CD8 T cells at the CR-B, CR-C, and −3.7 regulatory elements. The increased levels of these active histone marks correlated with the elevated levels of PD-1 expressed by LcKO CD8 T cells and mirrors the histone pattern observed in the PD-1 expressing cells at day 8 during chronic infection (Fig. 1C). Together, these results demonstrate that LSD1 is responsible for removing the activating histone marks H3K4^me1^ and H3K4^me2^ from the *Pdcd1* locus and suggest that LSD1 downregulates PD-1 expression by facilitating the removal of active histone marks after being recruited to locus by Blimp-1 during an acute viral infection.

### Epigenetic silencing of the locus following acute infection is enforced by LSD1

We previously showed that during the course of LCMV Armstrong infection antigen specific CD8 T cells initially lose CpG methylation at a region near CR-B (2). As the infection wanes and PD-1 levels return to near baseline, DNA methylation in that region reappears. Thus, loss of CpG methylation at the *Pdcd1* locus is correlated with gene expression, and reappearance of methylation is concurrent with gene silencing. Since *de novo* DNA methylation is dependent on an H3K4^me0^ methylation state (40, 41) and LSD1 has been shown to catalyze the removal of H3K4^me1^ and H3K4^me2^ (Fig. 5C), as well as being important for global DNA methylation (42), we sought to determine if remethylation of the *Pdcd1* was altered in LcKO mice during an acute infection. In accordance with the remethylation observed in WT animals at day 8 after LCMV Armstrong infection (2), WT control mice showed a relative abundance of CpG methylation across the *CR-B* region of *Pdcd1* (Fig. 6). In contrast, LSD1-deficient CD8 T cells, which have elevated levels of PD-1 expression at this time (Fig. 3), showed minimal remethylation of the locus with a majority of alleles analyzed exhibiting either no methylation or only a handful of methylated CpGs across the region (Fig. 6). This suggests that the inability to remove H3K4 methylation in LSD1-deficient CD8 T cells inhibits the acquisition of DNA methylation, which leads to a failure to fully silence PD-1, thereby providing a further mechanism for retaining prolonged gene expression.

**Figure 6.**
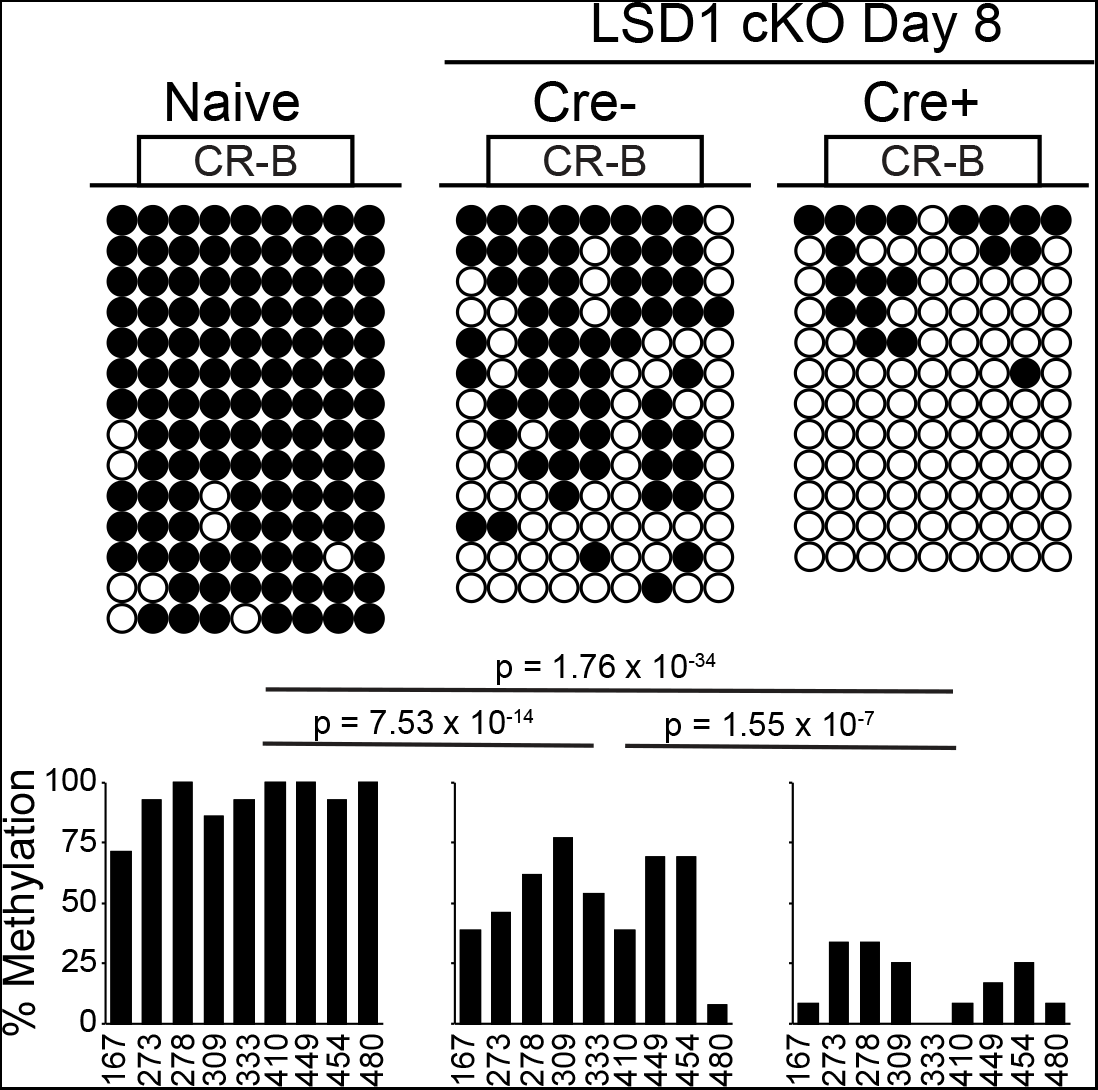
LSD1 deficient CD8 T cells fail to remethylate the *Pdcd1* locus during acute infection. LcKO and and *LSD1*^*fl/f*^*Cre*^−^ (WT) control mice were infected with LCMV Armstrong for 8 days. CD8 T cells from these mice or naïve, uninfected *Cre*- mice were isolated by MACS, and tetramer-specific cells were sorted by FACS. DNA from each population was bisulfite converted and PCR amplified. 8 clones from each mouse were sequenced, and incomplete sequences were discarded. At each CpG site across the CR-B region, the presence of DNA methylation within each clone is indicated by a closed circle and unmethylated CpG sites are indicated by an open circle. Frequency of methylation at each site across all clones is indicated in the corresponding bar graph. Data are combined from 2 biological replicates. Statistical significance was determined using a Fisher’s Exact Test and the P values are indicated.

### Increased frequency of PD-1 expressing CD8 T cells in the absence of LSD1 during melanoma

In order to investigate whether LSD1 deletion had an effect on PD-1 expression and disease progression in another model, we employed the B16 melanoma model (43). LcKO and *Cre*^−^ littermate controls were injected with 1×10^6^ tumor cells and mice were checked for weight and tumor size daily. Compared to each other, LcKO and WT mouse weights mice did not statistically differ throughout the course of the experiment; however, tumor growth appeared to be statistically lower in the mice with LSD-deficient CD8 T cells (Fig. 7A). Examination of the tumor infiltrating CD8+ T cells (TILs) showed that the mice which had LSD1-deficient cells had a higher frequency of PD-1^hi^ cells compared to control animals, suggesting that LSD1 deficiency plays a role in the expression of PD-1 in a non-viral inflammatory model (Fig. 7B).

**Figure 7.**
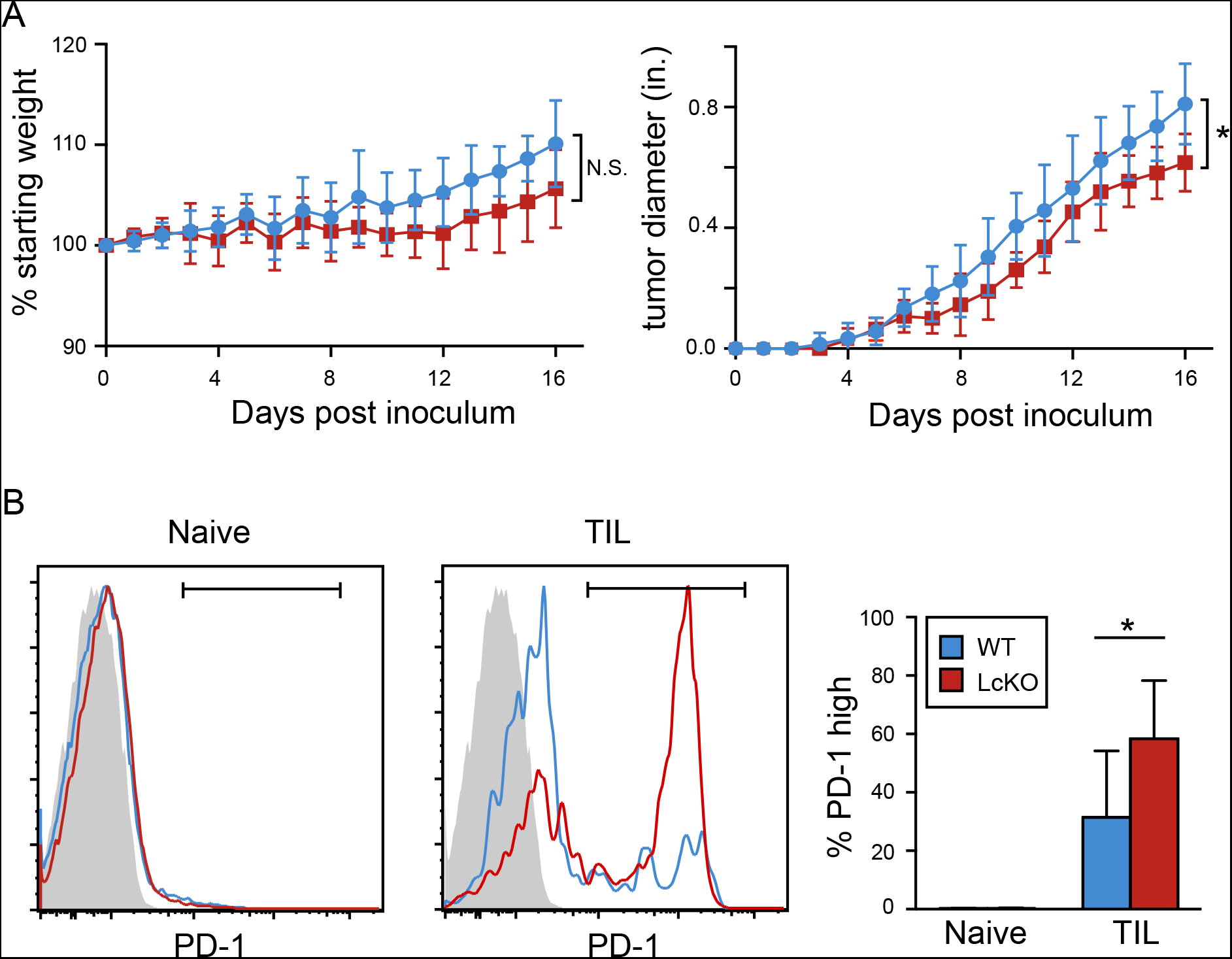
Percent PD-1 positive cells is increased in tumors from mice with LSD1-deficient CD8 T cells. LcKO and *LSD1*^*fl/f*^*Cre*^−^ (WT) control were injected with 1×10^6^ tumor cells and analyzed at days 16 or 17 post inoculation. (**A**) Percent starting weight and tumor diameter (inches) from mice during the 16 day time course. (**B**) At the end of the experiment, cells were collected from the spleen and tumor and subjected to flow cytometry. Activated CD8 T cells (CD44^hi^ CD62L ^low^) were gated on for their percentage of PD-1^hi^ cells in both the spleen and tumor (TIL). The summary graph in B shows the average of two independent experiments where the end points were day 16 and 17 post tumor cell inoculation. Significance was calculated in A using a two-way ANOVA (P = 0.015) and a Student’s *t* test in B. *, P < 0.05.

## Discussion

Multiple recent studies have highlighted the important effects of dynamic epigenetic regulation in driving immune responses (44–47). Here, we demonstrate that LSD1 is a novel epigenetic repressor of PD-1 expression. *Ex-vivo* stimulated LSD1-deficient CD8 T cells displayed increased levels of PD-1 expression compared to WT cells. LCMV-specific CD8 T cells also displayed increased PD-1 expression levels at day 8 following acute infection, but not chronic. Blimp-1 was identified as necessary to recruit LSD1 to *Pdcd1* regulatory regions using LSD1 and Blimp-1 conditional knockout mice. Although Blimp-1 associates with *Pdcd1* in both acute (PD-1 silencing) and chronic (PD-1 permissive) inflammatory environments, LSD-1 is only recruited to the locus in the acute infection, providing a mechanistic epigenetic switch for the differential expression of PD-1 in the two infectious systems. Furthermore, the association of LSD1 with *Pdcd1* during resolution of an acute infection correlates with and is necessary for the disappearance of the active histone modifications targeted by LSD1: H3K4^me1^ and H3K4^me2^. Additionally, the actions of LSD1 and the removal these histone marks correlate with the remethylation of the CR-B upstream, regulatory region and the silencing of PD-1 protein expression.

The results from this work also show that in a melanoma cancer model, the tumor infiltrating CD8 T cells from the LcKO mice displayed higher PD-1 levels compared to wild-type controls. This matches the observation seen in the acute viral infection where activated CD8 T cells deficient in LSD1 expressed higher levels of PD-1. Despite the increased presence of PD-1^hi^ cells, the overall health (as measured by weight loss) of the conditional knockout animals was not significantly different than in the wild-type mice. However, LcKO mice had smaller tumors across the time course. This could be due to LSD1 controlling other genes that also show enhanced expression when LSD1 is deleted in this model. In line with this, deletion of LSD1 in plasmablasts resulted in the upregulation of 471 genes compared to wild-type cells (28). Pharmacological inhibition of epigenetic modifiers, including LSD1 are effective in treating some cancers as a method to directly inhibit genes aberrantly expressed in the cancer itself (48). Thus, although the model used in our research does not affect LSD1 within the tumor cells, the higher PD-1 on TILs and potential subsequent cellular exhaustion could complicate this treatment strategy.

Blimp-1, a protein associated with B cell maturation into antibody-secreting plasma cells, is a critical inhibitor of *Pdcd1* transcription following acute T cell stimulation (19). Increases in Blimp-1 mRNA and protein following antigen clearance are associated with concurrent decreases in PD-1. Paradoxically, if antigen persists and stimulation through the TCR continues, Blimp-1 levels nonetheless increase further (21). Whereas in an acute infection the CD8 T cells from BcKO mice showed increased PD-1, suggesting an inhibitory function, the same deletion resulted in modestly lower levels of PD-1 in a chronic infection, suggesting Blimp-1 acted as an activator. In this study, Blimp-1 was shown to be bound at *Pdcd1* in both infection modalities, thereby precluding the possibility that the changing function of Blimp-1 was mediated indirectly through binding to other target genes. Instead, recruitment of LSD1, which was dependent on the presence of Blimp-1, was found to be unique to acute infections.

Many mechanisms could potentially explain the failure of Blimp-1 to recruit LSD1 in a chronic infection setting. Splice variants of Blimp-1 have been shown both to be involved in different timing of expression (49) and to have alternative functions (50). These include variants that alter Blimp-1’s ability to recruit additional transcription factors while preserving its ability to bind DNA (21, 51). Other biochemical mechanisms could also be involved, including posttranslational modifications of either Blimp-1 or LSD1. Furthermore, other factors binding to locus could sterically hinder or aid LSD1 recruitment. Another mechanism explaining the differential binding of LSD1 during acute and chronic infection could be expression levels of LSD1, however analysis of published transcriptomics data sets (39) suggest that this is not the case.

For *Pdcd1*, LSD1 provides an important mechanistic link between the dynamics of histone code modifications, DNA methylation, and gene expression of this critical immune regulatory locus. Dynamic molecular events at the *Pdcd1* locus are mirrored in chromatin accessibility patterns and DNA methylation patterns that change as CD8 T cells differentiate from naïve to effector, memory, and exhausted cells at both *Pdcd1*, as well as many other T cell expressed genes (2, 10, 15, 45–47). While this study focused on the effects of epigenetics on determining the expression of a single immune-related gene, LSD1 is most likely to have additional consequences on regulating CD8 T cell differentiation and gene expression. Irrespective of LSD1’s additional roles in modulating CD8 T cell gene expression, the work presented in this study establishes a clear mechanism for LSD1 to differentially regulate the expression levels of PD-1 during acute and chronic viral infections.

## Acknowledgments

We thank members of the lab for helpful critical and comments during the course of this work. We thank Royce Butler for expert animal husbandry. We thank Dr. Rafi Ahmed for providing LCMV viral strains; the National Institute of Health Tetramer Core Facility for providing LCMV specific tetramers; and also R. Karaffa and K. Fife for cell sorting from the Emory University School of Medicine Flow Cytometry Core.

## Abbreviations

Blimp-1: B lymphocyte induced maturation protein-1
ChIP: Chromatin immunoprecipitation assay
CR-B: Conserved region B
CR-C: Conserved region C
LCMV: Lymphocytic choriomeningitis virus
LSD1: Lysine specific demethylase 1
PD-1: Programmed cell death 1.

---

**Supplemental Table 1.**
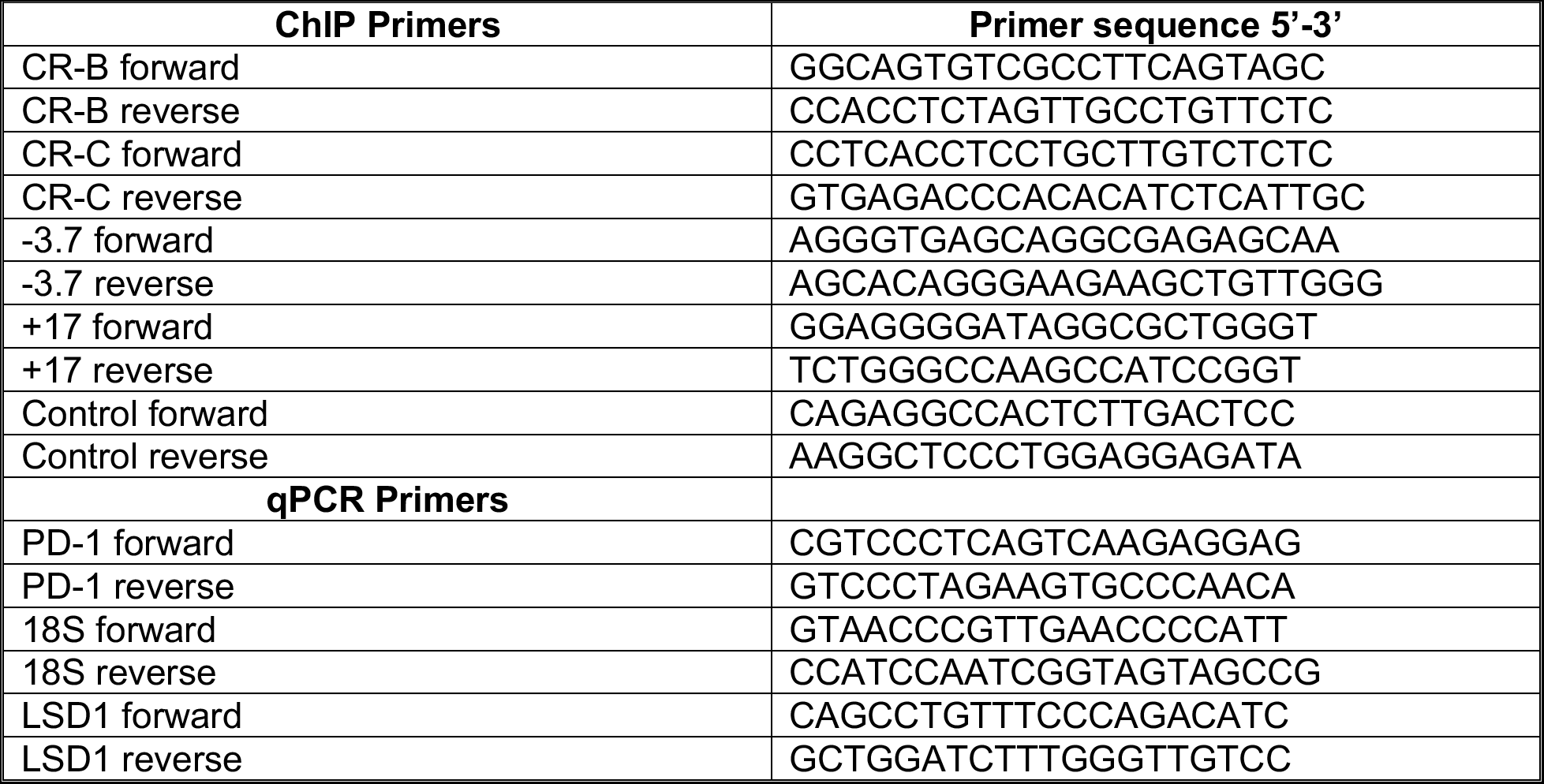
Primers used

